# Multi-target biocontrol by avocado fungal endophytes and their metabolites

**DOI:** 10.64898/2026.04.19.719497

**Authors:** Daniel Sánchez-Hernández, Luis A. Ibarra-Juárez, Cecilia I. Mayo-Montor, Juan L. Monribot-Villanueva, José A. Guerrero-Analco, John Larsen, Frédérique Reverchon

## Abstract

**Background:** Endophytic fungi can contribute to host defense against pests and pathogens through insecticidal and/or antifungal activity. Here, we evaluated the multi-target biocontrol potential of 11 avocado endophytic fungal strains against three plant pathogens (*Colletotrichum gloeosporioides*, *Fusarium solani* and *Phytophthora cinnamomi*) and two insect pests (*Sitophilus zeamais* and *Xyleborus bispinatus*) and analyzed the metabolomic profile of the two fungal strains with strongest bioactivity.

**Results:** *Trichoderma* species exhibited the strongest pathogen inhibition in dual-culture and diffusible compound assays. *Purpureocillium* and *Pochonia* strains combined pathogen inhibition through diffusible metabolites with strong insecticidal activity, supporting their multi-target potential. Ethyl acetate (EtOAc) extracts from strain *Trichoderma* 3 inhibited all phytopathogens, whilst strain *Purpureocillium* extracts were most active against *P. cinnamomi*. Compounds tentatively identified as macrocyclic trichothecenes (*Trichoderma* 3) and cyclosporine-like cyclic peptides (*Purpureocillium*) were potentially responsible for the displayed bioactivity.

**Conclusion:** Avocado endophytic fungi and their crude extracts show promising potential for multi-target biocontrol, whilst their metabolomic profiles provide candidate compounds for further chemical isolation and bioactivity evaluation.

## 1 INTRODUCTION

Avocado (*Persea americana* Mill.) is a globally important crop, with Mexico being the production and export world leader. In 2024, 253,044 hectares of avocado were cultivated in Mexico, for a total production of 2,761,327.85 tons.^1^ However, avocado production faces significant phytosanitary challenges, including insect pests and fungal diseases that negatively impact productivity and quality of the export product.^2^

Among the quarantine insect pests affecting avocado, members of the order *Coleoptera*, family *Curculionidae* —such as *Macrocopturus aguacatae*, *Conotrachelus* spp., and *Heilipus lauri*— are of particular concern. These pests are considered under official control, but their presence restricts domestic and international trade.^3,4^ Additionally, ambrosia beetles (*Coleoptera*: Curculionidae: Scolytinae), which bore into tree bark and cultivate symbiotic fungi as a food source, are regarded as major pests due to their destructive impact on avocado. Ambrosia beetles also act as vectors of phytopathogenic fungi such as *Fusarium* spp. or *Harringtonia lauricola,* hence further hampering avocado production.^5–8^ Their presence in Mexican avocado orchards has been recently recorded.^9^

Fungal and oomycete diseases are other limiting factors for avocado cultivation.^10^ Key phytopathogens include soil-borne microorganisms, such as *Phytophthora cinnamomi* and *Fusarium* spp. causal agents of root rot.^11,12^ Particularly, *P. cinnamomi* has been listed as one of the most damaging invasive species worldwide.^13^ Fruit pathogens, on the other hand, include *Sphaceloma perseae*, responsible for avocado scab*, as well as Colletotrichum gloeosporioides*, *Glomerella cingulata*, *Alternaria* spp., and *Dothiorella* spp.^14–16^ Among those, avocado anthracnose caused by *C. gloeosporioides*is the most relevant phytosanitary problem in post-harvest, inducing up to 50% loss of the fruit production.^17,18^ The above-mentioned pests and diseases significantly affect the health and productivity of avocado orchards, thus emphasizing the importance of integrated pest and disease management in avocado cultivation.^19^ Although management strategies for pest and pathogen mitigation heavily rely on chemical and cultural control,^20^ increasing regulatory restrictions on pesticide use, along with growing consumer demand for environmentally sustainable products, underscore the need for eco-friendly management strategies.^21^

One promising alternative to the use of pesticides is the application of beneficial endophytic microorganisms as biological control agents. Endophytic fungi inhabit plant tissues asymptomatically and produce a wide array of bioactive secondary metabolites with antimicrobial and insecticidal properties,^21–24^ highlighting their potential for biocontrol of pests and pathogens (hereafter referred as multi-target biocontrol). For example, spores and filtrates of *Trichoderma hamatum*, a root endophyte of *Brassica oleracea*, induced mortality of cotton leaf worm, possibly through the action of a siderophore.^25^ In another study, culture filtrates of endophytic *Beauveria bassiana* affected the development of *Helicoverva zea* larvae, beauvericin being identified as one of the potentially responsible metabolites.^26^ Other insecticidal compounds produced by known entomopathogenic fungi include destruxin A (DTX A) from *Metarhizium brunneum,* which increased mortality in *Bemisia tabaci* nymphs and *Spodoptera littoralis* larvae feeding on fungus-colonized melon and tomato leaves, respectively.^27,28^ These findings demonstrate the diversity of fungal compounds that could be used as bases for the development of bioinsecticides and call for further research on compounds produced by endophytic strains beyond the well-studied entomopathogenic *Metarhizium* and *Beauveria*.

The antifungal activity of endophytic fungal strains and their metabolites has also been documented. For example, the Chinese fir endophyte *Epicoccum dendrobii* and its organic extracts inhibit mycelial growth and spore germination of *C. gloeosporioides in vitro* and *in planta,* although the identity of antifungal metabolites was not investigated.^29^ Zhao *et al*.^30^ reported that griseofulvin, isolated from the root endophyte *Nigrospora* sp., exhibited broad-spectrum antifungal activity, significantly inhibiting the growth of eight important phytopathogens. More recently, Elias *et al*.^31^ reported the fungistatic activity of a polyformic acid and cytochalasin D, produced by an endophytic strain of *Xylaria*, against *C. gloeosporioides.* The anti-oomycete potential of endophytic fungi has also been described in dark septate root endophytes. For example, *Phialocephala europaea* was reported to inhibit the growth of *Phytophthora citricola,* most likely through the production of extracellular metabolites such as sclerin, sclerotinin A, and sclerotinin B.^32^ In avocado, research on antifungal activity displayed by fungal endophytes has been largely focused on the genus *Trichoderma*.^33–35^ Moreover, Nieves-Campos *et al*.^36^ showed that avocado fungal endophytes such as *Penicillium* sp., *Metapochonia* sp. and *Mortierella* sp. not only inhibited *P. cinnamomi in vitro* but also displayed plant growth-promoting effects.

The abovementioned studies have focused on a single-target approach, addressing either insect pests or phytopathogens. Although the multi-target biocontrol potential of entomopathogenic fungi isolated from insects has been recently investigated,^37^ the antifungal and insecticidal activity of endophytic fungal strains has been scarcely addressed. One exception is study by Safavi *et al*.^38^ who reported the multi-target biocontrol activity of two wheat endophytic strains of *Metarhizium anisopliae* against the red spider mite *Tetranychus urticae* and against the phytopathogenic fungus *Alternaria alternata* in tomato. Moreover, endophytic colonization by *M. anisopliae* stimulated tomato growth, possibly through the priming of plant defense responses, and highlight the potential of endophytic fungi as multi-target biocontrol agents.

Based on the evidence summarized above, we hypothesized that some of the avocado fungal endophytes isolated from our previous study^39^ may possess multi-target biocontrol potential against key pests and pathogens, and that this *in vitro* bioactivity is mediated, at least partly, by the metabolites they produce. We thus evaluated the multi-target biocontrol potential of 11 avocado endophytic strains against the phytopathogens *C. gloeosporioides*, *F. solani*, and *P. cinnamomi*, and against the insect pests *Sitophilus zeamais* and *Xyleborus bispinatus*. The chemical profiles of the two most promising strains were then analyzed through untargeted metabolomics to provide a chemical basis for their biocontrol performance.

## 2 MATERIALS AND METHODS

### 2.1 Biological material

Eleven fungal strains, previously isolated from the roots and branches of healthy avocado trees cultivated in Michoacán, Mexico,^39^ were selected based on their taxonomic identity and on previous reports of their bioactivity against pests and/or pathogens (**Table 1**).

**Table 1.** Endophytic fungal isolates selected in this study, their GenBank accession number, closest NCBI matches, and reported bioactivity.

| Endophytic fungal strain | GenBank accession number (NCBI) | Accession numbers of best matching sequences (NCBI) | Strain identifier in this work* | Bioactivity |  | Target organism | Reference |
|---|---|---|---|---|---|---|---|
|  |  |  |  | Antifungal / anti-oomycete | Insecticidal / Nematophagous |  |  |
| *Epicoccum aff. endophyticum* TCCA5-1 | PQ684548 | NR_172436.1 | *Epicoccum 1* | Inhibition of mycelial growth | --- | *OOMYCETES: Pythium irregulare*<br>*FUNGI: Fusarium graminearum* | 40, 41 |
| *Epicoccum aff. endophyticum* TCOA6-4 | PQ684612 | NR_172436.1 | *Epicoccum 2* |  |  |  |  |
| *Gaeumannomyces aff. hyphopodioides* TCCR3-2S | PQ684531 | NR_155138.1 | *Gaeumannomyces* | Induced host systemic resistance | --- | *FUNGI: Gaeumannomyces tritici* | 42 |
| *Pochonia aff. chlamydosporia* TNCR6-2S | PQ684629 | NR_171784.1 | *Pochonia 1* | --- | Nematophagous | *NEMATODES: Meloidogyne* spp.<br>*Pratylenchus*, *Radopholus* | 43,44 |
| *Pochonia aff. chlamydosporia* TCOR5-2S | --- | --- | *Pochonia 2* |  |  |  |  |
| *Purpureocillium aff. lavendulum* TCCR5-1S | PQ684512 | NR_166039.1 | *Purpureocillium* | --- | Production of bioactive compounds, parasitism | *NEMATODES: Meloidogyne incognita*<br>*INSECTS: Bemisia tabaci* | 45, 46 |
| *Talaromyces aff. assiutensis* UROR1-8 | PQ684505 | NR_172040.1 | *Talaromyces* | Inhibition of mycelial growth | Entomopathogenic fungi | *FUNGI: Sclerotinia sclerotiorum*<br>*INSECTS: Spodoptera litura, Chilo suppressalis, Ostrinia furnacalis, Rhopalosiphum padi, Acyrthosiphon pisum*, | 47 |
| *Trichoderma aff. longibrachiatum* URCA2-4 | PQ684730 | NR_120298.1 | *Trichoderma 1* | Inhibition of mycelial growth, mycoparasitism, plant growth promotion | Entomopathogenic fungi | *FUNGI: Botrytis, Colletotrichum, Erysiphe, Fusarium, Magnaporthe, Sclerotinia, Verticillium, Rhizoctonia,*<br>*OOMYCETES: Pythium, Phytophthora*<br>*INSECTS: Spodoptera littoralis* | 25, 48, 49 |
| *Trichoderma aff. spirale* URCR1-5 | PQ684706 | NR_077177.1 | *Trichoderma 2* |  |  |  |  |
| *Trichoderma aff. spirale* URCR6-3S | PQ684701 | NR_077177.1 | *Trichoderma 3* |  |  |  |  |
| *Trichoderma sp* TCCA2-5 | --- | --- | *Trichoderma 4* |  |  |  |  |

\* Strain identificator in this work = shorter strain name to simplify the terminology in the manuscript.

The antifungal and anti-oomycete activity of these fungal strains was evaluated against three economically important avocado pathogens: *C. gloeosporioides*, *F. solani*, and *P. cinnamomi*. Strains of *C. gloeosporioides* (described in Xoca-Orozco *et al*.^50^) and *P. cinnamomi* (strain TGR1-5) were provided by Julio Vega (Universidad Nacional Autónoma de México) and Sylvia Fernández-Pavia (Universidad Michoacana San Nicolás de Hidalgo), respectively. Strain *F. solani* B1 was isolated from an avocado orchard in Veracruz, Mexico.^12^

Insecticidal activity of the endophytic fungal strains was evaluated against two agriculturally important pest species: *S. zeamais* and *X. bispinatus. Sitophilus zeamais* is a cosmopolitan species that affects stored grains and serves as a model to evaluate insecticidal activity due to its easy maintenance in laboratory conditions, its rapid life cycle and its susceptibility to entomopathogenic fungi.^51^ In contrast, *X. bispinatus* is a ambrosia beetle with increasing relevance in avocado agroecosystems as it may act as a vector of avocado fungal pathogens.^52^ Prior to their experimental use, insects were raised and fed under controlled conditions to ensure their physiological homogeneity.^53^

### 2.2 *In vitro* antagonism assays against phytopathogens

The antagonistic activity of the selected endophytic fungi was evaluated through three *in vitro* approaches: dual culture confrontation, double sealed plate assays to assess the antifungal/anti-oomycete activity of volatile organic compounds (VOCs), and antagonistic activity of diffusible metabolites. All experiments were conducted with three replicates per treatment.

#### 2.2.1 Dual culture confrontation

Dual culture confrontation was performed as described by Fokkema^54^ and Nieves-Campos *et al*.,^36^ in Petri dishes containing potato dextrose agar (PDA). A 5-mm diameter mycelial plug of the phytopathogen was placed on one edge of the plate, while a plug of the endophytic fungal strain was positioned on the opposite edge. Plates were incubated at 27°C for 14 days. At the end of the incubation period, the mycelial growth of phytopathogens was documented through photographic records and measured using ImageJ software (version 1.54p). The percentage of growth inhibition (PGI) was calculated as follows:

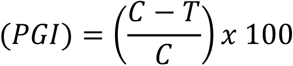

where ***C*** is the diameter growth of the pathogen in the control plate and ***T*** is the diameter growth in the presence of the endophytic fungus.

#### 2.2.2 Double sealed plate assay

The inhibitory effect of fungal VOCs was evaluated against the three phytopathogens as described by Meena *et al*.^55^ Five-mm mycelial plugs of the endophytic fungus and the phytopathogen were placed at the center of separate Petri dishes containing PDA. The two plates were placed face-to-face and sealed with Parafilm®. Control treatments consisted of the same system with the phytopathogen only. Plates were incubated at 27°C for seven days, after which the growth of the phytopathogen was photographed and colony diameters were measured using ImageJ. The PGI was calculated as described above.

#### 2.2.3 Diffusible compound assay

The inhibitory effect of diffusible compounds secreted by endophytic fungi was evaluated as described by Meena *et al*.^55^ Sterile microporous cellophane membranes were placed over the surface of PDA plates. A 5-mm mycelial plug of the endophytic fungus was inoculated onto the membrane and incubated at 27°C until it fully colonized the surface. The cellophane membrane with the fungal biomass was then removed. Immediately afterward, a 5-mm mycelial plug of the phytopathogen was placed in the center of the plate to grow in contact with the diffused metabolites. Control plates consisted of PDA without exposure to endophytic fungi. After incubation at 27°C for seven days, diameter growth of the phytopathogen was measured and the PGI was calculated as described previously.

### 2.3 Insecticidal assay by spore suspension dipping

The insecticidal activity of the 11 endophytic fungal strains was evaluated against *S. zeamais* and *X. bispinatus* through direct-contact bioassays using spore suspension dipping, as described by Carrillo *et al*.^56^ Each fungal strain was cultured on PDA for 10 to 14 days at 27°C to promote conidial production. Spore suspensions were prepared by adding sterile distilled water to the surface of the fungal cultures and gently scraping with a sterile loop. The resulting suspensions were filtered through sterile gauze and adjusted to a final concentration of 1 × 10⁸ conidia ml-1, except for the two strains of *Epicoccum*, for which the maximum concentration that could be reached was 1 × 106 conidia ml-1.

For each fungal strain, 30 homogeneous-age adult insects of each species were randomly selected and divided into three replicates (10 insects per replicate). Individual insects were immersed into 1.5 ml of the spore suspension for 90 seconds. After treatment, insects were transferred to sterile Petri dishes containing moistened filter paper. Negative controls were immersed into sterile distilled water only. A positive control treatment with a 1:50 cypermethrin solution (commercial insecticide, ANAJALSA®) was also included. All experimental units were maintained at 26°C with a 12:12 h light–dark photoperiod. Insect survival and mortality were recorded daily for 7 to 13 days, depending on insect survival. Dead insects were not replaced but removed after mortality confirmation. Direct observations were made using a stereoscopic microscope to verify fungal colonization emerging from insect bodies.

Daily mortality data were used to construct survival curves and cumulative mortality trajectories. Mortality data were expressed as percentages and corrected using Abbott’s formula^57^:

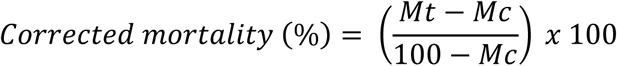

where ***Mt*** is the mortality percentage in the treated group, and ***Mc*** the mortality percentage in the control group.

### 2.4 Liquid culture and extraction

Two fungal strains were selected, based on their multi-target biocontrol potential: *Trichoderma aff. spirale* URCR6-3S (hereafter *Trichoderma* 3) and *Purpureocillium aff. lavendulum* TCCR5-1S (*Purpureocillium*). Erlenmeyer flasks with 300 ml of potato dextrose broth (PDB) were inoculated with 15 mycelial plugs each (5-mm diameter), excised from seven-day-old PDA cultures and incubated at 27°C for nine days under continuous agitation (200 rpm). Each strain was cultured in triplicate, and PDB was used as a control. The culture broth was then centrifuged at 6,000 rpm for 20 min at 4°C. The resulting cell-free supernatants were transferred to glass bottles previously washed with sterile distilled water and methanol to minimize contamination prior to diffusible compounds extraction.

Culture supernatants were subjected to sequential liquid–liquid extraction with organic solvents using a 1:1 (v/v) solvent-to-supernatant, following Guevara-Avendaño *et al*.^58^. The supernatant was first extracted three times with ethyl acetate (EtOAc) in a separatory funnel by vigorous shaking. After each extraction, the organic phase was recovered and pooled, while the remaining aqueous phase was retained and extracted three times with n-butanol (*n*-BuOH), following the same procedure. The same extraction procedure was applied to uninoculated PDB (control). The organic extracts were dried over anhydrous sodium sulfate and concentrated under reduced pressure using a BÜCHI RII rotary evaporator at 40°C. Fungal extracts obtained from each solvent were then weighed to determine extraction yield, transferred to clean glass vials, and stored at −20°C for subsequent bioassays.

### 2.5 Antifungal activity of organic extracts

The antagonistic activity of EtOAc and *n*-BuOH extracts was evaluated against *C. gloeosporioides*, *F. solani*, and *P. cinnamomi* according to Guevara-Avendaño *et al*.^59^. Each phytopathogen was cultured in PDB by transferring a mycelial fragment from an active PDA culture and incubating it at 25°C for 24 h with constant shaking (200 rpm). After incubation, 1 ml of the liquid culture was added to 100 ml of molten PDA, homogenized, and poured into sterile Petri dishes. Once solidified, a central hole was made in each plate using a cork borer, and 100 µl of crude extract solution (5 mg ml⁻¹ in methanol) were added to each hole. Assays were performed in triplicate for each phytopathogen and crude extract obtained from *Purpureocillium* and *Trichoderma* 3. Thiabendazole was included as a positive control, while methanol was used as a solvent control. Inhibition halos were measured after incubation at 27°C for 72 h.

### 2.6 Metabolomic analysis

Untargeted metabolic profiling was performed with ultra-high-performance liquid chromatograph ACQUITY UPLC I-Class coupled to a quadrupole time-of-flight (Synapt G2Si, HDMi) mass spectrometer (Waters, Milford, MA, USA). Chromatographic separation was carried out on an ACQUITY BEH column (1.7 μm, 2.1 x 50 mm, C18; Waters, USA) with a column and sample temperatures of 40°C and 15°C, respectively. Mobile phase consisted of (A) water and (B) acetonitrile, both containing 0.1% formic acid (LC-MS grade; Sigma-Aldrich). Gradient conditions of the mobile phases were 0-20 min linear gradient 1-99% B, 20-24 min 99% B under isocratic conditions, 24-25 min linear gradient 99-1% B (total run time was 30 min). Flow rate was 0.3 ml min-1 and 5 µl of the extract at 20 mg ml-1 was injected. Mass spectrometric analysis was performed with an electrospray ionization source in positive and negative modes with a capillary, sampling cone and source offset voltages of 3000, 40 and 80 V, respectively. Source temperature was 120°C and desolvation temperature was 150°C. Desolvation gas flow was 650 l h-1 and nebulizer pressure was 6.0 bar. Leucine-enkephalin was used as the lock mass (556.2771, [M+H]+; 554.2615, [M-H]-). Method for mass acquisition was MS^e^ and conditions were: 50-1200 *m/z* mass range, collision energy for function 1 was 6 V, and for function 2, a ramp from 10 to 30 V. Scan time was 0.5 s. Data were acquired with MassLynx v4.1 (Waters, USA).

Raw mass spectrometry data were processed using Progenesis QI v3.0 (Nonlinear Dynamics, Newcastle, UK), including peak detection, retention-time alignment, spectral deconvolution, and generation of a feature matrix defined by the corresponding *m/z* and retention-time values. Putative metabolite identification was performed by comparing experimental *m/z* values and fragmentation patterns of detected features with simulated MS/MS spectra of compounds included in the LOTUS Natural Products database (https://lotus.naturalproducts.net/), available through PubChem (https://pubchem.ncbi.nlm.nih.gov/), restricted to metabolites reported for Hypocreales; and with experimental reference MS/MS spectra from MS-DIAL mass spectral libraries (https://systemsomicslab.github.io/compms/msdial/main). Candidate annotations were selected based on an absolute mass error of ≤ 10 ppm, agreement between experimental and reference fragmentation patterns, and isotopic pattern similarity.

### 2.7 Statistical analyses

Data from antagonism assays were analyzed separately for each phytopathogen. Evaluation of variance between replicates and distribution were done using the Shapiro-Wilk normality test. When complete growth saturation occurred, as observed for *C. gloeosporioides* in some assays, treatment effects were assessed using the non-parametric Kruskal-Wallis test, followed by Dunn’s *post hoc* comparisons. In contrast, datasets without saturation (*F. solani* and *P. cinnamomi*) showed normal or near-normal distributions but heteroscedasticity, thus differences between treatments were evaluated using Welch’s one-way ANOVA with Games-Howell *post hoc* tests.

Survival and mortality data obtained from the insect bioassays were analyzed separately for each insect species. Daily survival data were used to construct survival curves for each fungal treatment and control using Kaplan-Meier estimator. Differences in survival distributions among treatments were evaluated using the log-rank test. Following a significant overall effect, pairwise log-rank comparisons were conducted with Holm correction for multiple testing. Cumulative mortality at the end of the bioassays was calculated for each treatment. Final mortality percentages were calculated for each replicate and corrected using Abbott’s formula when mortality in the negative control exceeded zero. Differences among treatments were evaluated using a Kruskal–Wallis rank sum test. All statistical analyses were performed in R^60^ using the *stats*, *FSA* and *rstatix* packages, as appropriate.

To integrate antagonistic performance against phytopathogens with insecticidal assays, scatter plots were constructed using scaled response variables. For each assay, inhibition percentage and Abbott-corrected mortality were normalized independently using min-max rescaling:

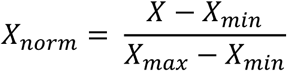

Were ***X*** representing the observed value within each assay.

A multi-target performance index was then calculated for each strain as the arithmetic mean of normalized inhibition and normalized mortality:

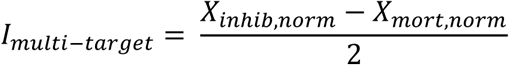

The antifungal activity of crude extracts was analyzed using one-way ANOVA. Residual normality and homogeneity of variances were assessed using Shapiro– Wilk and median-centered Levene’s tests. Each extract was compared with the corresponding methanol control using Dunnett’s test with adjusted *p*-values.

Metabolomic data (*m/z*_retention time values) obtained from both ionization modes were analyzed separately for each solvent using MetaboAnalyst 6.0 (https://www.metaboanalyst.ca/). Features with zero intensity across all samples were removed, and only those detected in at least two of the three biological replicates were retained. Data were normalized by the sample median, log₁₀- transformed, and Pareto-scaled. Global differences among the uninoculated PDB control, *Trichoderma* 3, and *Purpureocillium* were explored using principal component analysis (PCA), and their statistical significance was assessed by PERMANOVA based on 199 permutations.

Differential analysis was performed independently for EtOAc and *n*-BuOH extracts using moderated *t*-statistics implemented in limma (MetaboAnalyst 6.0). Three pairwise comparisons were conducted within each extract: *Purpureocillium* versus *Trichoderma* 3, *Purpureocillium* versus uninoculated PDB, and *Trichoderma* 3 versus uninoculated PDB. *P*-values were adjusted for multiple testing using the Benjamini–Hochberg false discovery rate (FDR). Features with FDR < 0.05 and |log₂FC| ≥ 1, corresponding to a fold change ≥ 2 or ≤ 0.5, were considered differential.

Strain-associated features (significantly enriched in one strain relative to the other strain and to the control) were identified by intersecting the significant feature lists based on their unique identifiers. Because of the larger number of differential features in the EtOAc extracts, its list intersections were performed in RStudio. Hierarchical clustering of the retained EtOAc candidates was conducted using Euclidean distance and Ward’s linkage, and relative abundance patterns were visualized as row-wise z-scores.

## 3 RESULTS

### 3.1 Pathogen inhibition by avocado fungal endophytes in dual culture confrontation

For the dual-culture assays against *C. gloeosporioides,* three fungal strains showed significant differences compared to the control (Kruskal-Wallis, H = 34.12, df = 11, p < 0.001), all belonging to the *Trichoderma* genus. *Trichoderma* 1 exhibited the highest percentage of inhibition (64.72%), followed by *Trichoderma* 3 (42.64%) and *Trichoderma* 2 (37.48%).

Fungal endophytes with significant growth inhibition of *F. solani* were *Gaeumannomyces* (67.12%), followed by *Trichoderma* 3 (63.23%) and *Trichoderma* 4 (49.77%) (Welch’s ANOVA, *F* (11, 8.76) = 37.90, *p* < 0.001). Although *Trichoderma* 1 and *Trichoderma* 2 showed high inhibition values (75.16% and 53.82%, respectively), no significant differences were detected with the control due to variability between replicates.

All endophytic strains significantly inhibited the mycelial growth of *P. cinnamomi* (Welch ANOVA, *F* (11,9.15) = 598.76, *p* < 0.001). The highest inhibitory activity (> 65%) was displayed by strains of *Trichoderma* and *Gaeumannomyces*, while the remaining strains exhibited inhibition percentages ranging from 26.02% to 56.30% (**Figure 1A**).

**Figure 1.**
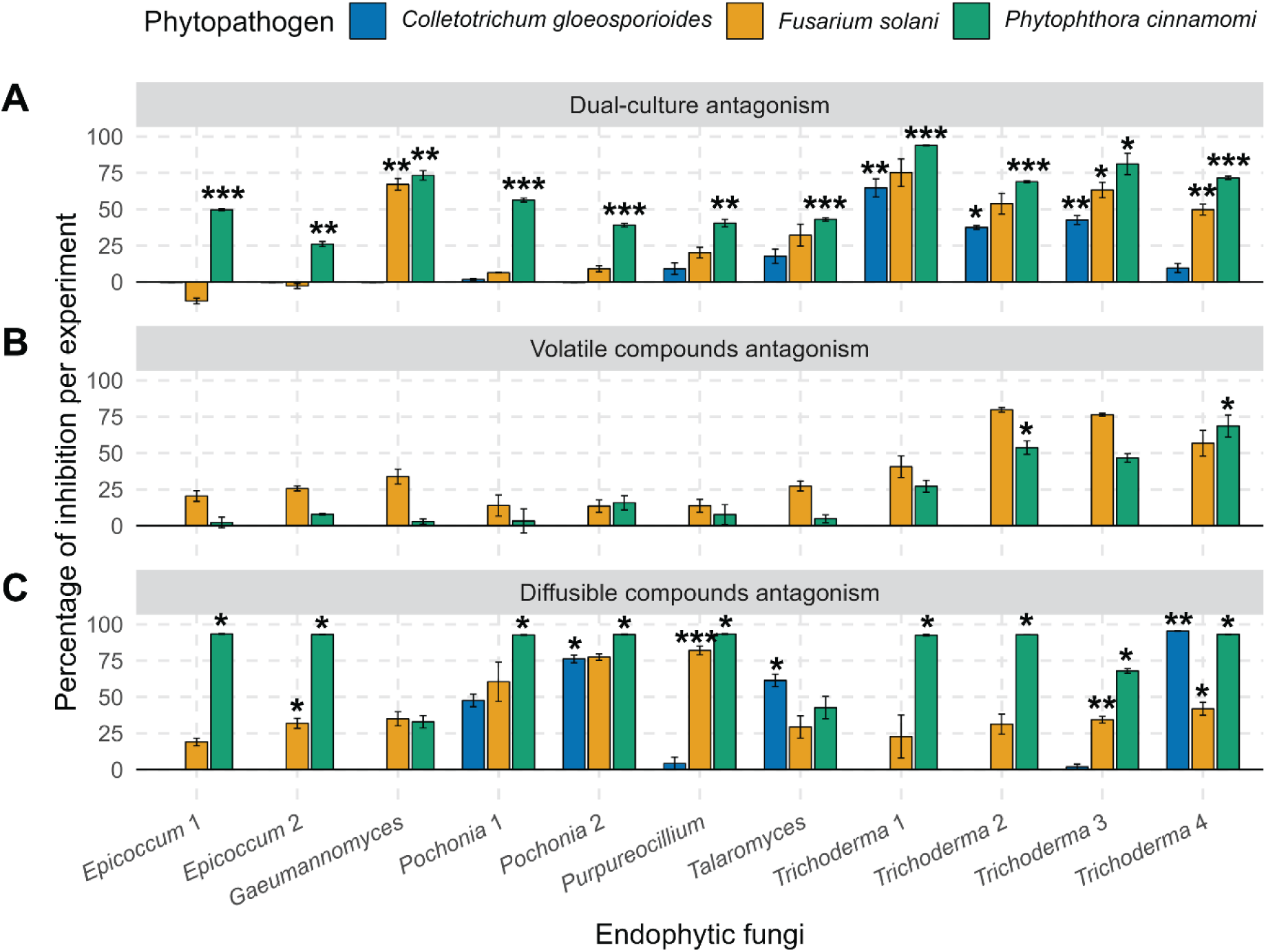
Antagonistic activity of endophytic fungi against avocado phytopathogens. Percentage of inhibition per experiment against *C. gloeosporioides*, *F. solani*, and *P. cinnamomi* evaluated through dual-culture (A), volatile compounds (B) and diffusible compound (C) antagonistic assays. Bars represent mean ± standard error (SE) (n=3). Statistical analyses were performed separately for each phytopathogen following diagnostic evaluation of data distribution and variance. Asterisks indicate significant differences between each individual treatment and the control (* = *p* < 0.05, ** = *p* < 0.01, *** = *p* < 0.001), based on Dunn’s *post hoc* test following Kruskal-Wallis analyses or Games-Howell *post hoc* test following Welch’s one-way ANOVA, as appropriate.

### 3.2 Pathogen inhibition through VOC emission

None of the eleven endophytic fungi evaluated in this study showed inhibitory capacity against *C. gloeosporioides* through VOC emission. For *F. solani*, Welch’s ANOVA showed significant differences between treatments (*F* (11,9.6) = 72.26, *p* < 0.001); however, *post hoc* comparisons using the Games-Howell test did not detect significant differences between the control and treatments (*p* (adj) > 0.05), even though some strains of *Trichoderma* showed high inhibition percentages (> 55%). The large confidence intervals suggest that data variability limited the detection of significant differences at the strain level. In antagonistic assays with *P. cinnamomi*, Welch’s ANOVA showed statistically significant differences between treatments (*F* (11,9.01) = 19.14, *p* < 0.001) and only in two individual treatments compared to the control: *Trichoderma* 4 (56.83% inhibition) and *Trichoderma* 2 (53.77%) (**Figure 1B**).

### 3.3 Pathogen antagonism through diffusible compounds

In antagonism assays via diffusible metabolites against *C. gloeosporioides*, three endophytic strains showed significant differences compared to the control: *Trichoderma* 4 (95.41%), followed by *Pochonia* 2 (76.15%) and *Talaromyces* (61.38%) (Kruskal-Wallis analysis, H = 32.66, df = 11, p < 0.001). Welch’s ANOVA combined with the Games-Howell *post hoc* analysis showed that *Purpureocillium* (82.01%), *Pochonia* 2 (77.43%), *Trichoderma* 4 (41.89%) and *Epicoccum 2* (31.91%) displayed significant inhibition against *F. solani* (*F* (11,9.38) = 55.96, *p* < 0.001). Finally, nine endophytic fungal strains were able to reduce the growth of *P. cinnamomi* through diffusible compounds, with inhibition percentages greater than 90% in eight strains (Welch’s ANOVA and Games-Howell *post-hoc* test, *F* (11,9.05) = 51.97, *p* < 0.001). In contrast, *Gaeumannomyces* and *Talaromyces* did not show significant differences compared to the control (**Figure 1C**).

### 3.4 Insecticidal assays against *Sitophilus zeamais*

Kaplan-Meier analysis revealed significant differences in *S. zeamais* survival distributions between fungal treatments (log-rank test, *p* < 0.01). The cypermethrin positive control caused rapid and almost complete insect mortality, as only one insect survived the immersion during the 12 days of the trial. Among fungal treatments, *Purpureocillium* produced the most potent insecticidal effects, showing a marked and progressive reduction in survival over time. Pairwise logarithmic rank comparisons with Holm correction confirmed that only the *Purpureocillium* treatment differed significantly from the negative control (*p* < 0.001). Other fungal strains exhibited little effect on insect survival, with curves partially overlapping with that of the negative control (**Figure 2A**).

**Figure 2.**
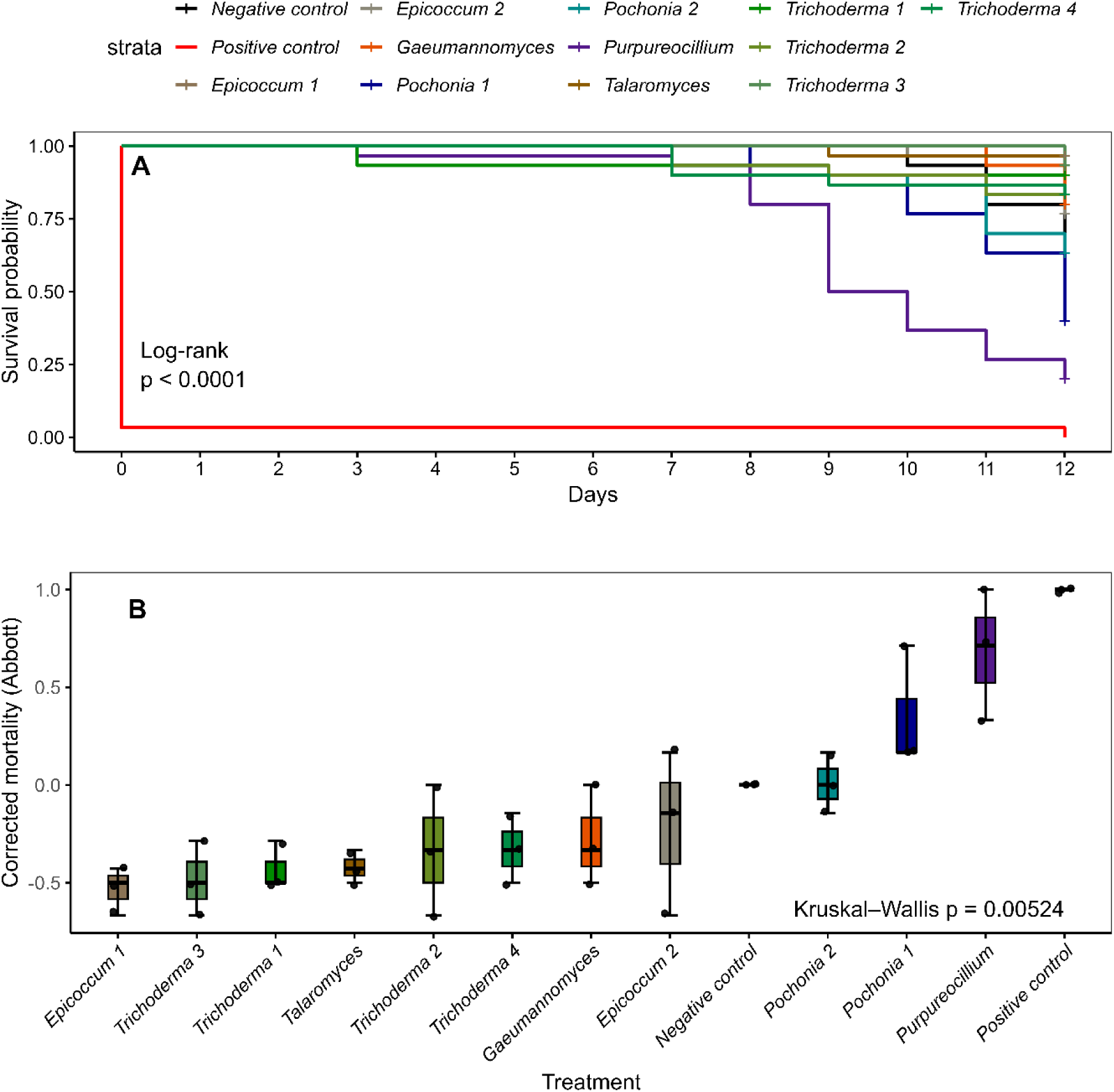
Kaplan-Meier survival curves and Abbott-corrected final mortality of *Sitophilus zeamais* following exposure to fungal treatments. A) Survival probabilities were estimated over a 12-day period. The positive control (commercial insecticide) caused rapid mortality, while fungal treatments showed variable effects on insect survival. B) Abbott-corrected final mortality presented as boxplots. Values represent correct mortality percentages calculated per replicate (n=10).

Abbott-corrected final mortality supported the survival analysis, with *Purpureocillium* (68.42%) and *Pochonia* 1 (36.84%) showing the highest corrected mortality values, while the remaining fungal treatments resulted in lower mortality levels (**Figure 2B**).

### 3.5 Insecticidal assays against *Xyleborus bispinatus*

Survival curves for *X. bispinatus* differed significantly between treatments according to Kaplan-Meir analysis (log-rank test, *p* < 0.01). Rapid mortality at the start of the bioassay (five minutes after dipping) was observed for the positive control, which remained clearly separated from the fungal treatments and the negative control over time. *Pochonia 1* and *Purpureocillium* produced the strongest reductions in survival, showing a sustained and progressive effect compared to the negative control. Pairwise log-rank tests with Holm correction supported these observations (*p* < 0.01 for both treatments), indicating significant differences between these treatments and the negative control, as well as with the less effective fungal strains (**Figure 3A**). Moderate differences in survival were observed for *Pochonia* 2, *Trichoderma* 1 and *Trichoderma* 2, although not significantly when compared with the negative control. The remaining fungal treatments showed survival trajectories that largely overlapped with the negative control.

**Figure 3.**
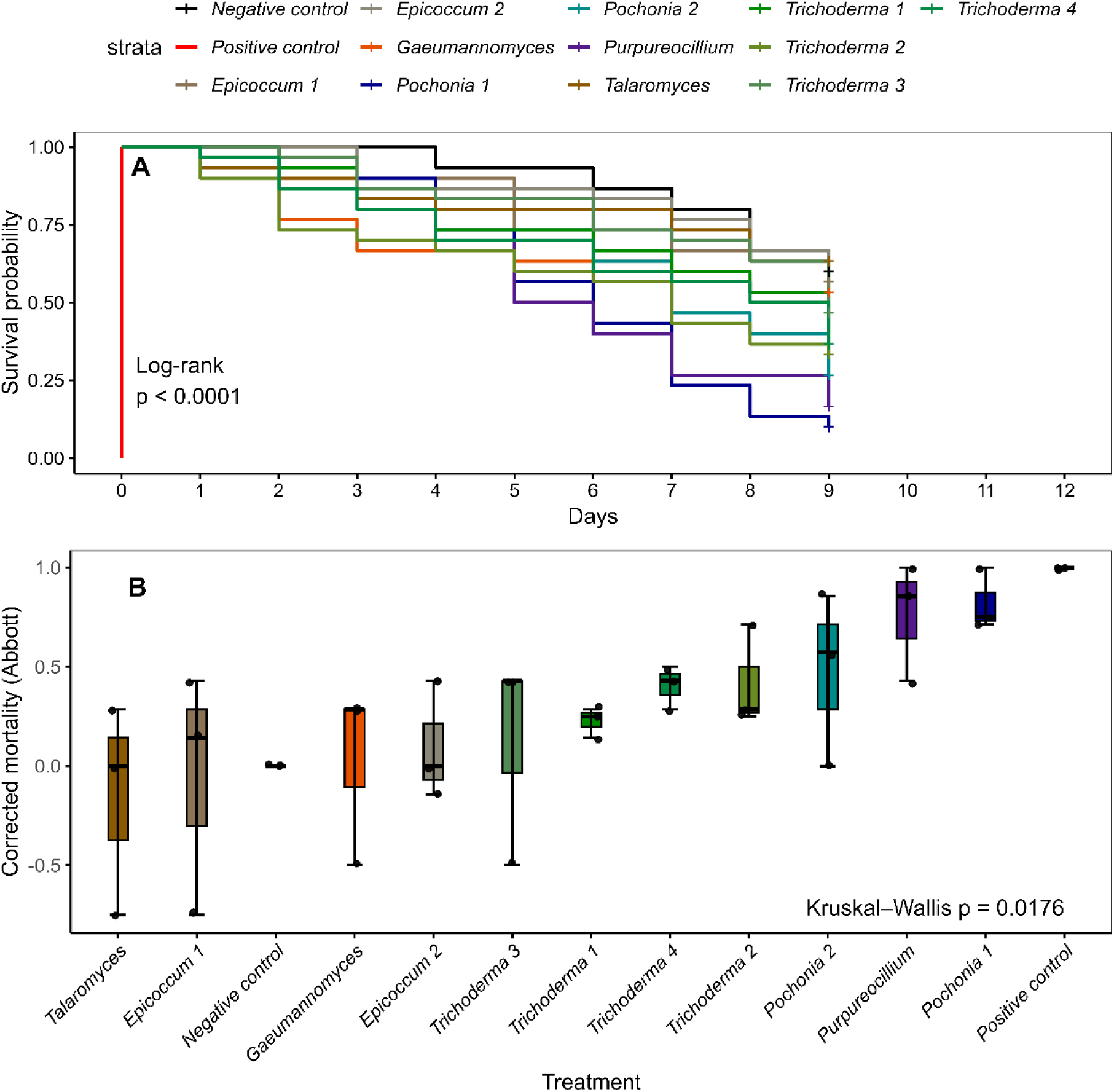
Kaplan-Meier survival curves and Abbott-corrected final mortality of *Xyleborus bispinatus* following exposure to fungal treatments. A) Survival probabilities were estimated over a 9-day period. The positive control (commercial insecticide) caused rapid mortality, while fungal treatments showed variable effects on insect survival. B) Abbott-corrected final mortality presented as boxplots. Values represent correct mortality percentages calculated per replicate (n=10).

The Abbott-corrected final mortality patterns reflected the survival results, highlighting *Pochonia* 1 (83.33 %) and *Purpureocillium* (72.22 %) as the most effective treatments against *X. bispinatus* (**Figure 3B**).

### 3.6 Multi-target biocontrol potential of endophytic strains

The scatter plot, which integrates the pathogen inhibition through three mechanisms and corrected insect mortality by spore suspension, revealed heterogeneous performance among the endophytic strains. Most treatments were concentrated at low Abbott-corrected mortality values, even though they exhibited moderate to high levels of pathogen growth inhibition, suggesting a limited correlation between antimicrobial and insecticidal activity (**Figure 4**). Variable results from dual-culture assays and the general lack of effects of fungal VOCs resulted in no fungal strain clustering in the multi-target upper right quadrant that combined pathogen growth inhibition with insecticidal activity. However, when diffusible compound antagonism and insect mortality were considered, *Purpureocillium* and *Pochonia* 1 were highlighted for their multi-target biocontrol activity (**Figure 4**).

**Figure 4.**
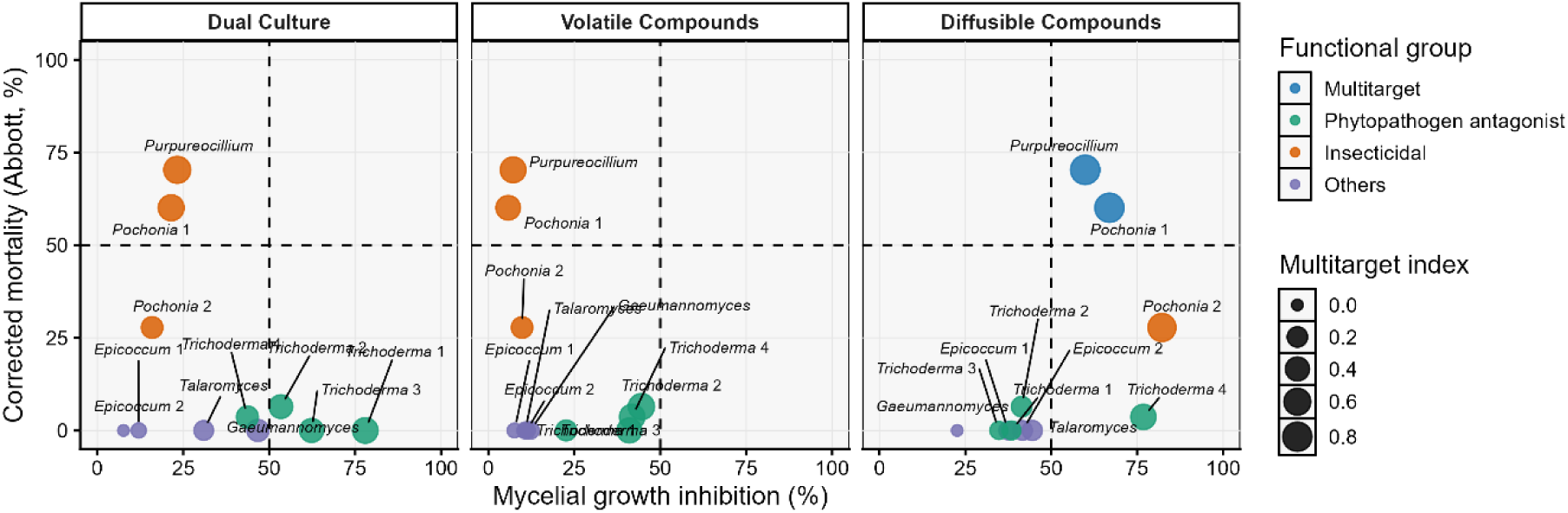
Scatter plot integrating the antifungal and anti-oomycete performance and insecticidal activity of endophytic fungal strains. Each point represents the scaled response of a treatment and assay combination, where the inhibition of mycelial growth against three phytopathogens, combined via dual culture, volatile, and diffusible metabolite assays is plotted against Abbott-corrected insect mortality obtained from exposure to the spore suspension.

### 3.7 Inhibitory activity of fungal crude extracts against pathogens

Based on its multi-target biocontrol potential, *Purpureocillium* was selected for chemical profiling. Moreover, *Trichoderma* 3 was added to the metabolic profiling analysis as a contrasting strain, since it displayed strong antifungal activity but weak insecticidal activity. Crude-extract yields are reported in **Supplementary Material 1**.

The EtOAc extract of *Trichoderma* 3 produced significantly larger inhibition halos than the methanol control against all pathogens. The remaining extracts did not differ significantly from the control against *C. gloeosporioides* or *F. solani*. However, all extracts produced significantly larger halos than the control against *P. cinnamomi*. No inhibition halos were observed in the methanol controls (**Figure 5**).

**Figure 5.**
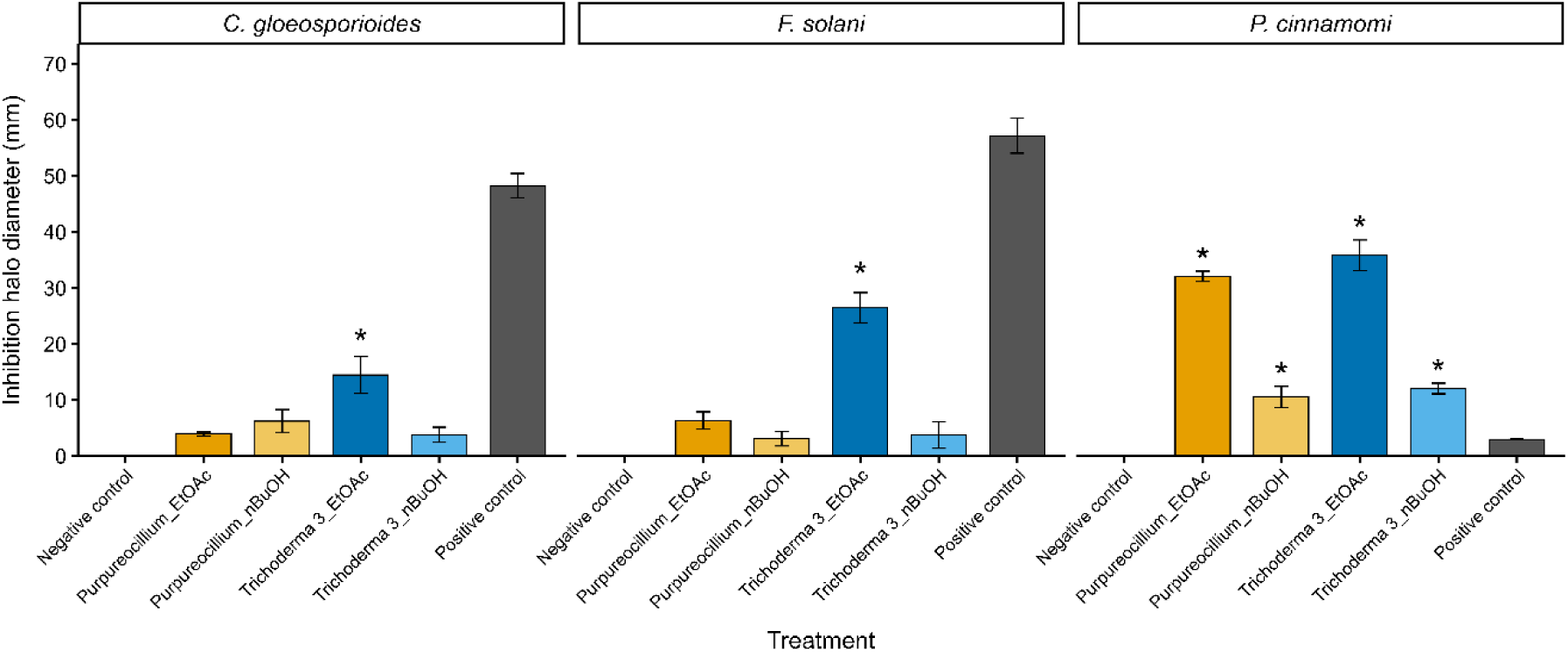
Inhibitory activity of metabolite extracts from *Purpureocillium* and *Trichoderma* 3 against *C. gloeosporioides*, *F. solani*, and *P. cinnamomi*. Extracts were obtained using ethyl acetate (EtOAc) and *n*-butanol (*n*-BuOH). Bars represent the mean inhibition halo diameter (mm) ± standard error (n = 3). Methanol and thiabendazole served as the negative and positive controls, respectively. Asterisks indicate significant differences relative to the negative control (*p* < 0.05). The positive control is shown solely as a reference and was excluded from statistical comparisons.

### 3.8 Untargeted metabolomic analysis

Untargeted metabolomic profiling was performed independently for the EtOAc and *n*-BuOH extracts because of their different physicochemical selectivity. Following data filtering, 215 and 55 compounds (*m/z*_Rt pairs values) were retained in the EtOAc and *n*-BuOH datasets, with each dataset encompassing samples from both fungal strains and the PDB control. PCA revealed distinct metabolic patterns among *Purpureocillium, Trichoderma* 3 and the control in both solvent extracts. In the EtOAc extract, PC1 and PC2 explained 57.4% and 15.3% of the total variance, respectively (72.7% combined). *Trichoderma* 3 was clearly separated from the PDB control and *Purpureocillium* along PC1, whereas the control and *Purpureocillium* were mainly differentiated along PC2. Replicates of *Trichoderma* 3 and the control clustered closely, while *Purpureocillium* showed greater within-group variability. In the *n*-BuOH extract, PC1 and PC2 explained 30.1% and 21.4% of the total variance (51.5% combined). PERMANOVA based on the PC1–PC2 scores indicated significant differences among the metabolomic profiles of the PDB control, *Purpureocillium*, and *Trichoderma* 3 in both solvent extracts. Group identity explained 96.6% of the variation in the EtOAc extract (*F*-value = 86.19, *R*² = 0.966, *p* = 0.020) and 50.9% in the *n*-BuOH extract (*F*-value = 3.11, *R*² = 0.509, *p* = 0.015) (**Supplementary Material 2**).

### 3.9 Metabolic profiling and selection of strain-associated compounds

Differential analysis of the EtOAc extract identified 118 compounds that differed between *Purpureocillium* and *Trichoderma* 3 (FDR < 0.05 and |log₂FC| ≥ 1). Of these, 51 were enriched in *Purpureocillium* and 67 in *Trichoderma* 3. Comparison with the PDB control identified nine differential compounds for *Purpureocillium*: eight enriched in the fungal culture and one in PDB. For *Trichoderma* 3, 126 compounds differed from PDB, including 57 enriched in the fungal culture. Intersection of the significant compound lists using their unique identifiers retained seven compounds enriched in *Purpureocillium* relative to both *Trichoderma* 3 and PDB, and 49 compounds enriched in *Trichoderma* 3 relative to both *Purpureocillium* and PDB. Thus, 56 compounds from the EtOAc extracts were retained as strain-associated metabolites (**Supplementary Material 3**).

In the *n*-BuOH extract, two compounds differed between *Purpureocillium* and *Trichoderma* 3. Of these, only one was significantly enriched in *Purpureocillium* relative to PDB and therefore retained as strain-associated. This compound was tentatively annotated as indole-3-acetamide (PubChem CID: 397).

Differential abundance patterns between fungal cultures were visualized using volcano plots for both solvent extracts (**Figures 6A-B**). A heatmap with hierarchical clustering was generated for the 20 highest-ranked compounds in the EtOAc extract, grouping biological replicates according to treatment and revealing distinct sets of compounds enriched in *Trichoderma* 3 and *Purpureocillium* (**Figure 6C**). Heatmap visualization was restricted to the EtOAc extract since only one strain-associated compound was retained from the *n*-BuOH extract.

**Figure 6.**
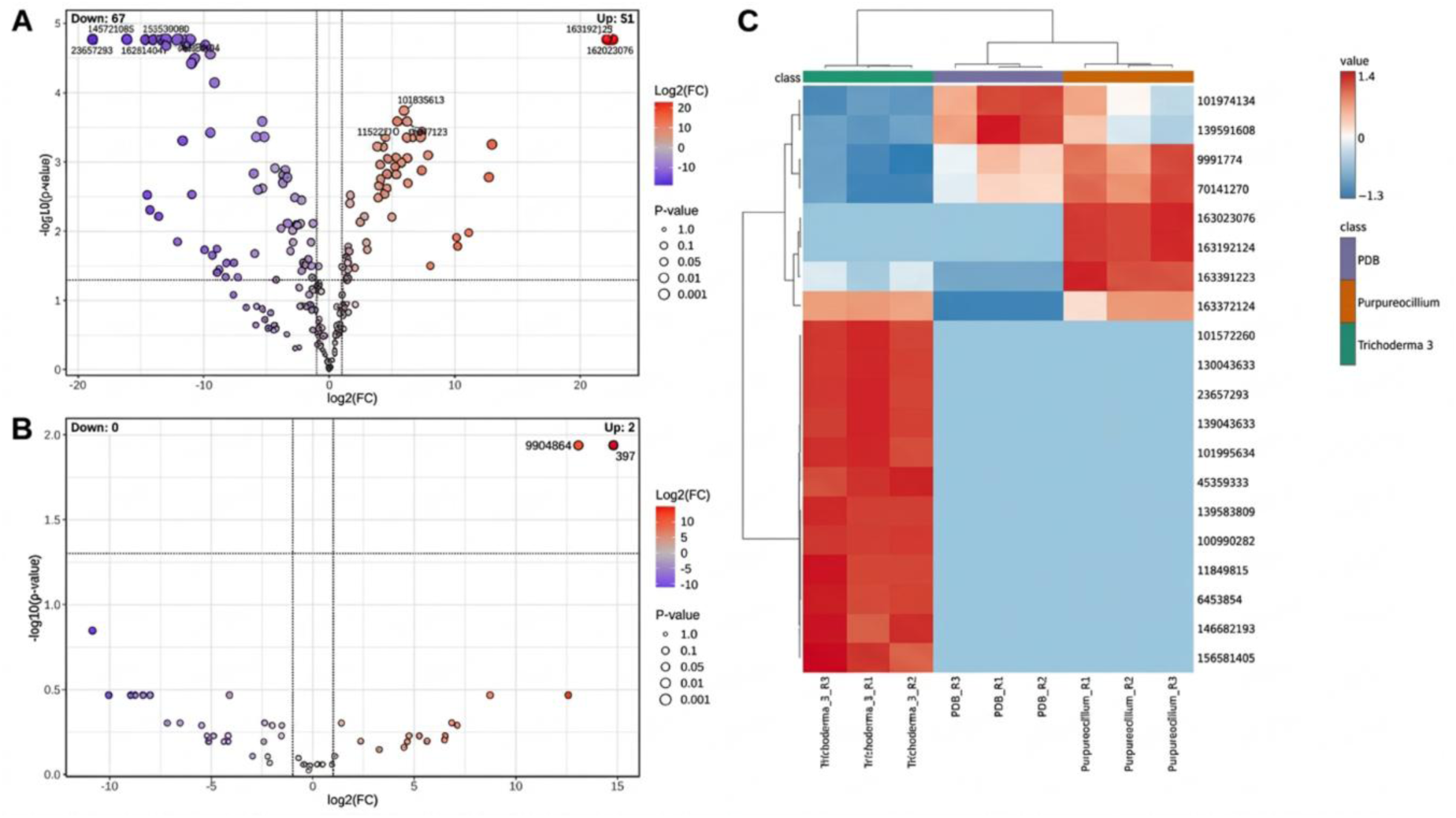
Differential and hierarchical clustering analyses of strain-associated compounds in EtOAc and *n*-BuOH extracts. (A–B) Volcano plots showing differential compounds between *Purpureocillium* and *Trichoderma* 3 in the (A) EtOAc and (B) *n*-BuOH extracts. Each point represents an individual compound. Positive log₂FC values indicate enrichment in *Purpureocillium*, whereas negative values indicate enrichment in *Trichoderma* 3. Vertical and horizontal dotted lines represent the thresholds |log₂FC| ≥ 1 and FDR < 0.05, respectively. The EtOAc extract contained 51 compounds enriched in *Purpureocillium* and 67 in *Trichoderma* 3, whereas the *n*- BuOH extract contained two compounds enriched in *Purpureocillium* and none in *Trichoderma* 3. (C) Hierarchical clustering heatmap of the 20 highest-ranked strain-associated compounds retained from the EtOAc extract. Colors represent row-standardized relative abundances, with red and blue indicating higher and lower abundances, respectively. Samples and compounds were clustered using Euclidean distance and Ward’s linkage. Biological replicates grouped according to treatment, revealing contrasting abundance patterns among *Trichoderma* 3, *Purpureocillium*, and the uninoculated PDB control.

### 3.10 Tentatively annotated strain-associated metabolites in fungal extracts

Among the strain-associated compounds, 22 candidates from the EtOAc extract and one from the *n*-BuOH extract were tentatively annotated and selected for concise presentation based on their potential biological relevance (**Supplementary Material 4**). Selected compounds included cyclosporin P-like and cyclosporin C-like compounds associated with *Purpureocillium*, and macrocyclic trichothecenes associated with *Trichoderma* 3. Their chemical structures are presented in **Figures 7** and **8**.

**Figure 7.**
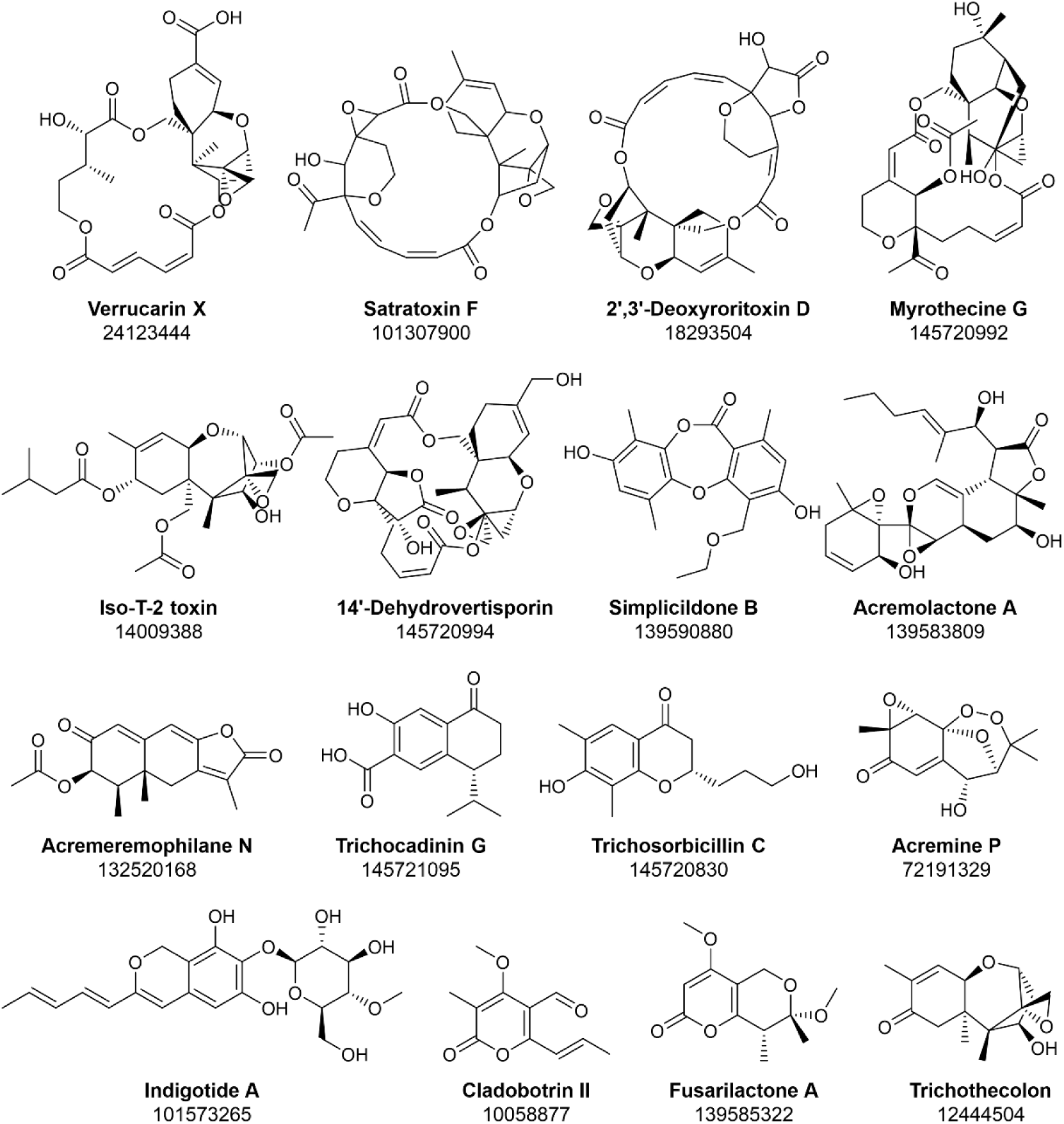
Chemical structures of selected putatively-annotated metabolites in EtOAc extracts from *Trichoderma* 3. Compound names and PubChem CIDs are provided below each structure.

**Figure 8.**
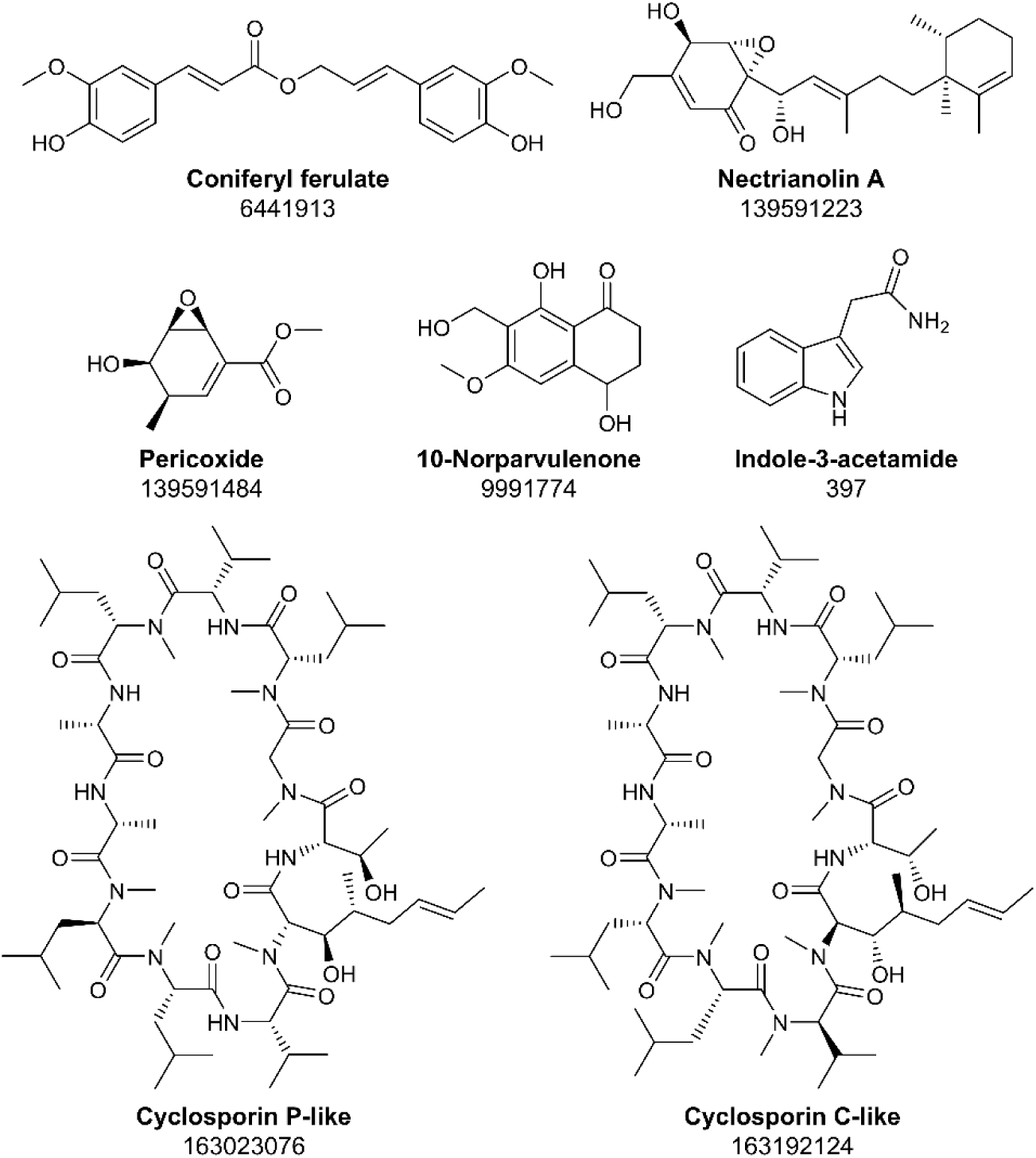
Chemical structures of selected putatively-annotated metabolites in EtOAc and *n*-BuOH extracts from *Purpureocillium*. Compound names and PubChem CIDs are provided below each structure.

## 4 DISCUSSION

Our results showed that the endophytic fungi included in this study differed in their antifungal activity, both in terms of spectrum and efficacy. Strains of *Trichoderma* showed high inhibition percentages of phytopathogens, with values exceeding 80% via dual culture and diffusible metabolites, although VOCs were generally less effective. On the other hand, only a subset of endophytic fungal strains exhibited insecticidal effects against *S. zeamais* and *X. bispinatus*, particularly those belonging to the genera *Purpureocillium* and *Pochonia*. Accordingly, the multi-target index distinguished these genera for their combined antifungal and insecticidal performance, highlighting the functional diversity and complementary biocontrol potential of the evaluated avocado endophytes.

Strains of *Trichoderma* have been widely reported as strong antagonists of crop pathogens through mycoparasitism, competition for space and resources, and production of antibiotic metabolites.^91,92^ In avocado, Hakizimana *et al*.^33^ and Andrade-Hoyos *et al*.^34^ documented the antagonistic activity of endophytic *Trichoderma* spp. against *P*. *cinnamomi*. Furthermore, López-López *et al*.^35^ described inhibitory effects of *Trichoderma* spp., isolated from avocado roots, against *P. cinnamomi* and *C. gloeosporioides*. In our study, *C. gloeosporioides* showed greater tolerance to fungal exposure and metabolites produced by endophytic fungi than *F. solani* and *P. cinnamomi*. The fact that endophytic strains with the greatest inhibitory capacity against *C. gloeosporioides* belonged to the genus *Trichoderma* reaffirms its importance for the biocontrol of avocado pathogens.

The high representation of macrocyclic trichothecene annotations in the EtOAc extract of *Trichoderma* 3, including verrucarin X, satratoxin F, 14′- dehydrovertisporin, or 2′,3′-deoxyroritoxin D, sheds some light on the possible mechanisms underlying its antifungal activity. The tentative identification of macrocyclic trichothecenes is also consistent with the presence of trichothecene biosynthetic gene clusters in *Trichoderma* species^93^ and the structural diversity of this toxin family.^71^ Macrocyclic trichothecenes inhibit protein synthesis in eukaryotic cells^99^ and exhibit antifungal and cytotoxic activities.^69^ Verrucarin X, for example, has been reported as a highly cytotoxic compound that preferentially inhibits protein biosynthesis *in vivo*.^64^ Macrocyclic trichothecenes from the fungal phytopathogen *Myrothecium roridum* also display antifungal activity against other plant pathogens.^65,66^ These reports support the strong antifungal activity exhibited by *Trichoderma* 3 and its EtOAc extracts, although the tentative detection of T-2-toxin, a well-characterized mycotoxin,^68^ warrants further characterization of this strain toxicological profile prior its use for field application.

Beyond the putative trichothecenes, *Trichoderma* 3 extracts contained several metabolites with reported antifungal or antimicrobial properties. Cladobotrin II is an antifungal *α*-pyrone aldehyde produced by the mycoparasitic fungus *Cladobotryum varium*.^61^ Fusarilactone A belongs to a family of *β*-lactones reported in *F. solani* that inhibit the tea pathogen *Pestalotiopsis theae*.^63^ Trichocadinins are cadinane sesquiterpenes with antibacterial and antifungal activities,^73^ whilst the biosynthesis of indigotides, molecules originally isolated from *Cordyceps indigotica*79, was associated with a secondary-metabolite supercluster in *Metarhizium.*^80^ Collectively, the coexistence of multiple compound families with complementary biological targets is consistent with the strong antimicrobial arsenal displayed by *Trichoderma* strains.

The topical application of spores from *Trichoderma* spp. has been shown to display insecticidal activity against various orders of economically important insect pests, including: Hemiptera (*Schizaphis graminum, Bemisia tabaci*, *Cimex hemipterus*);^95-97^ Diptera (*Delia radicum*);^98^ Lepidoptera (*Spodoptera littoralis*, *Leucinodes orbonalis*)^99,25^ and Coleoptera (*Xylotrechus arvicola*, *Acanthoscelides octetus* and *Sitophilus oryzae*).^100,101^ However, in our study, the immersion of *S. zeamais* and *X. bispinatus* in spores of *Trichoderma* spp. had no insecticidal effect. This discrepancy may be due to the origin of the fungal strain, as in most of the studies cited above, *Trichoderma* strains were isolated either from insects or soils. Moreover, distinct experimental setups, such as fungal spore concentration, life stage of the insects or rearing conditions, may account for these contrasting results. Collectively, these findings point out that *Trichoderma* avocado endophytes should not be discarded as potential pest biocontrol agents, and their insecticidal activity should be further tested under other experimental conditions.

The two fungal genera that stood out for their insecticidal and, hence, multi-target activity were *Purpureocillium* and *Pochonia*. Most studies on *Pochonia* have focused on its nematophagous activity.^43,44^ However, it has also been demonstrated that *Pochonia chlamydosporia* can inhibit the growth of *Fusarium oxysporum* f. sp. *cubense* and reduce its colonization in banana plants under controlled conditions, confirming its antagonistic capacity beyond nematode parasitism.^102^ Recently, Rincy *et al*.^103^ showed that *P. chlamydosporia* could reduce the growth of several fungal and oomycete phytopathogens. Specialized metabolites such as bisevertinol, enniatin I and enniatin H produced by *P. chlamydosporia* are active against *Pythium myriotylum* and *Phytophthora* sp., and against the plant-parasitic nematode *Radopholus similis*, thereby confirming the anti-oomycete and nematicidal properties of *Pochonia*. Our findings indicate that the two *Pochonia* strains tested in this study exhibited antifungal and anti-oomycete activity, in particular through the production of diffusible compounds. We report here for the first time the ability of *Pochonia* strains to cause mortality in *S. zeamais* and *X. bispinatus*. As strain *Pochonia 1* showed higher levels of insect mortality than *Pochonia 2*, whilst the latter displayed a broader antagonistic spectrum against phytopathogens, our findings suggest a strain-specific bioactivity.

The endophytic fungal strain with strongest insecticidal activity against both pests was *Purpureocillium*. This genus is widely known for its ability to parasitize nematodes of agricultural importance.^45^ Furthermore, *Purpureocillium* spp. have been reported as entomopathogens of several pests. Taleb *et al*.^104^ described their activity against *Culex pipiens,* a mosquito transmitting viral diseases. *Purpureocillium* spp. have also been shown to display insecticidal activity against the tobacco aphid *Myzus persicae*, the mite *Tetranychus urticae,* the fall armyworm *S. frugiperda,*^105,106^ and against insects of the family Curculionidae,^107,108^ consistent with our results. The UPLC-MS/MS profiling of *Purpureocillium* EtOAc extract revealed two signals tentatively assigned as cyclosporin P-like and cyclosporin C-like compounds, which are cyclic non-ribosomal undecapeptides with reported insecticidal activity.^88^ Cyclosporin-like metabolites may have contributed to the insecticidal effects observed against *S. zeamais* and *X. bispinatus*, although their identity and activity should be confirmed with the use of commercial standards.

The activity of *Purpureocillium* spp. as antagonists of phytopathogens has not been extensively studied. Wang *et al*.^109^ identified 20 genes in *Purpureocillium lilacinum* involved in the synthesis of leucinostatins A and B and demonstrated the anti-oomycete effect of these two lipopeptides against *Phytophthora infestans* and *Phytophthora capsici*. Here, we show that *Purpureocillium* could inhibit the growth of *F. solani* and *P. cinnamomi* through the action of diffusible metabolites, and that its EtOAc and *n*-BuOH extracts significantly reduced the mycelial growth of *P. cinnamomi*. The tentative identification of coniferyl ferulate in the EtOAc extract of *Purpureocillium* may partly explain its antifungal activity, as coniferyl ferulate has been described as antimicrobial and antioxidant.^83,84^ The *n*-BuOH extracts of *Purpureocillium* contained indole-3-acetamide (IAM), a direct precursor of the auxin indole-3-acetic acid in the tryptophan-dependent IAM pathway,^110,90^ suggesting that *Purpureocillium* could potentially influence plant growth in addition to its multi-target biocontrol activity.

## 5 CONCLUSIONS

Our study highlights *Purpureocillium and Pochonia* as avocado endophytes with biocontrol properties against multiple target organisms, and *Trichoderma* as the genus displaying the strongest antifungal activity. Metabolomic characterization of *Trichoderma* 3 and *Purpureocillium* linked their bioactivity to tentatively identified compounds such as macrocyclic trichothecenes and cyclosporine-like cyclic peptides, respectively. Our findings thus provide a chemical basis for future research aimed at bioactivity-guided purification, compound identity confirmation and evaluation. These investigations —combined with biosafety studies and the validation of strains and their metabolites in greenhouse and field trials— will help determine the viability of avocado endophytic fungal strains as tools for sustainable management of avocado pests and diseases.

## ACKNOWLEDGEMENTS

D.S.H. acknowledges the SECIHTI postgraduate scholarship awarded to him. Thanks are extended to Isabel Moreno de Palma and Mayté Meneses Marín for their support with the insecticidal activity assays, and to Alfonso Méndez-Bravo for his help with figure edition. Furthermore, the authors thank Carolina Camacho-Vázquez and Erika Valencia-Mejía for their assistance in the fungal extract preparation. This research was funded by a SECIHTI-PRONACES-PEE grant (project “PERSEA”, number 322772).

## DATA AVAILABILITY STATEMENT

The data that support the findings of this study are available from the corresponding author upon reasonable request.

