## Supplementary Material 1 for "Multi-target biocontrol by avocado fungal endophytes and their metabolites"

|  | **EtOAc extract yield**  **(mg l^-1^)** | ***n*-BuOH extract yield**  **(mg l^-1^)** |
| --- | --- | --- |
| *Purpureocillium* strain | 76.8 ± 13.0 | 3,217.6 ± 431.3 |
| *Trichoderma* 3 strain | 267.2 ± 91.6 | 2,327.4 ± 702.5 |
| PDB control | 70.3 | 1,726.7 |

**Supplementary Material 1.** Yields of EtOAc and *n*-BuOH extracts from the endophytic fungal strains *Purpureocillium* and *Trichoderma* 3, and the potato dextrose broth (PDB) negative control. Extracts were obtained in triplicate for each strain, except for the PDB control which was only extracted once in each solvent.
