## Supplementary Material 2 for "Multi-target biocontrol by avocado fungal endophytes and their metabolites"

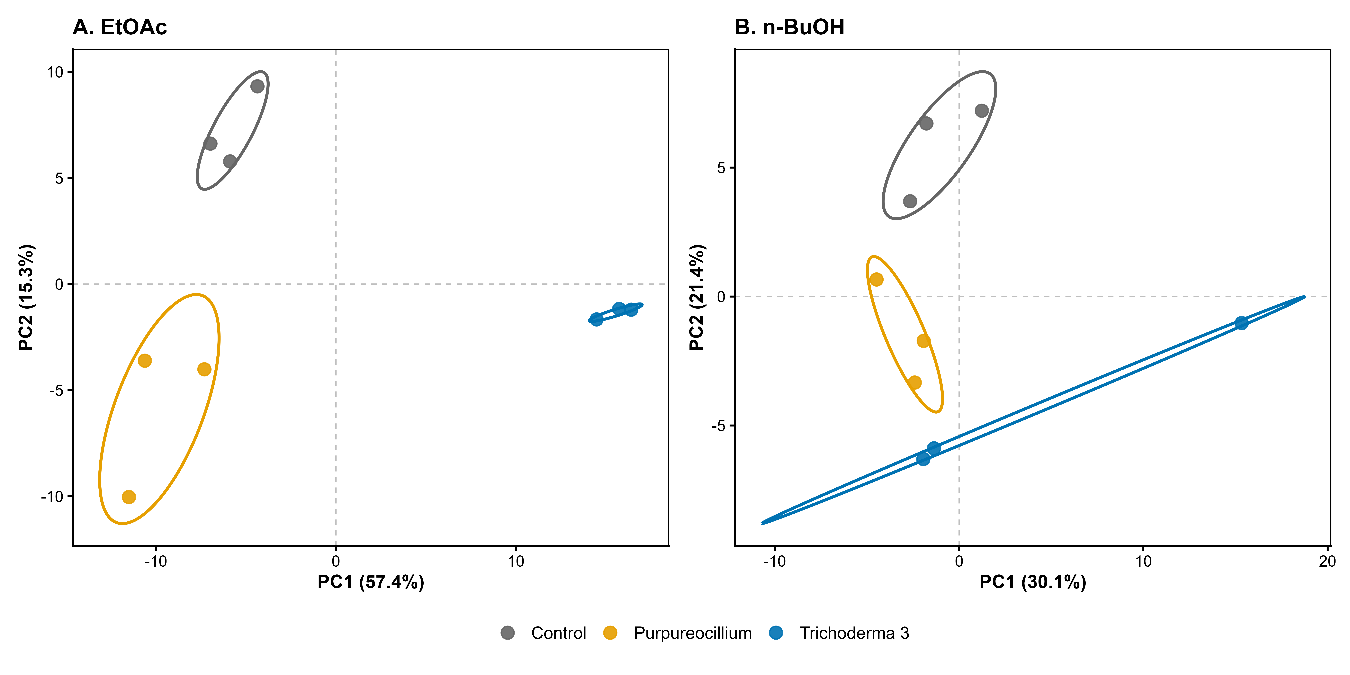
**Supplementary Material 2.** Principal component analysis (PCA) showing the distribution of metabolomic profiles among fungal strains and the uninoculated culture medium control based on EtOAc (A) and *n*-BuOH (B) extracts.
