## Supplementary Material 4 for "Multi-target biocontrol by avocado fungal endophytes and their metabolites"

| **PubChem ID** | **Putatively annotated metabolite** | **Strain/extract** | **Reported biological activity** | **Reference** |
| --- | --- | --- | --- | --- |
| 10058877 | Cladobotrin II | *Trichoderma* 3 / EtOAc | Antifungal; antibacterial activity (α-pyrone aldehyde reported in *Cladobotryum* *varium*) | 61,62 |
| 139585322 | Fusarilactone A | *Trichoderma* 3 / EtOAc | Antifungal activity; β-lactones from *Fusarium* solani inhibit *Pestalotiopsis theae* | 63 |
| 24123444 | Verrucarin X | *Trichoderma* 3 / EtOAc | Moderate antifungal and cytotoxic activities; inhibition of protein biosynthesis; zoosporicidal activity | 64,65 |
| 101307900 | Satratoxin F | *Trichoderma* 3 / EtOAc | Antiviral, anticancer, antimalarial, and antifungal activities | 66 |
| 14009388 | Iso-T-2 toxin | *Trichoderma* 3 / EtOAc | Inhibition of protein synthesis, oxidative stress induction, and apoptosis | 67,68 |
| 18293504 | 2',3'-Deoxyroritoxin D | *Trichoderma* 3 / EtOAc | Antifungal and cytotoxic activities (roritoxin family) | 69 |
| 145720994 | 14'-dehydrovertisporin | *Trichoderma* 3 / EtOAc | Antifungal, antiviral, antimalarial, and anticancer activities (compound family) | 70 |
| 12444504 | Trichothecolon | *Trichoderma* 3 / EtOAc | Mycotoxin; inhibitor of protein synthesis in eukaryotic cells | 71 |
| 145720992 | Myrothecine G | *Trichoderma* 3 / EtOAc | Cytotoxic macrocyclic trichothecene (compound family) | 72 |
| 145721095 | Trichocadinin G | *Trichoderma* 3 / EtOAc | Antibacterial and antifungal activities (trichocadinins B–G family) | 73 |
| 145720830 | Trichosorbicillin C | *Trichoderma* 3 / EtOAc | Antioxidant, anti-HIV, and antimicrobial activities (sorbicillinoid class) | 74, 75 |
| 72191329 | Acremine P | *Trichoderma* 3 / EtOAc | Inhibition of sporangial germination in *Plasmopara* *viticola* (compound family) | 76 |
| 139583809 | Acremolactone A | *Trichoderma* 3 / EtOAc | Herbicidal activity; congeners B and C inhibit seedling growth | 77 |
| 132520168 | Acremeremophilane N | *Trichoderma* 3 / EtOAc | Anti-inflammatory activity | 78 |
| 101573265 | Indigotide A | *Trichoderma* 3 / EtOAc | Antibacterial biosynthetic supercluster in *Metarhizium robertsii* | 79, 80 |
| 139590880 | Simplicildone B | *Trichoderma* 3 / EtOAc | Antifungal activity against *Cryptococcus neoformans* | 81 |
| 6441913 | Coniferyl ferulate | *Purpureocillium* / EtOAc | Antimicrobial, anticancer, antioxidant, and vasodilatory activities | 82, 83, 84 |
| 139591484 | Pericoxide | *Purpureocillium* / EtOAc | No specific activity reported; the genus *Periconia* produces bioactive metabolites | 85 |
| 139591223 | Nectrianolin A | *Purpureocillium* / EtOAc | Cytotoxic activity against HL-60 and HeLa cells | 86 |
| 9991774 | 10-Norparvulenone | *Purpureocillium* / EtOAc | Antiviral activity | 87 |
| 163023076 | Cyclosporin P-like | *Purpureocillium* / EtOAc | Insecticidal activity reported for several members of the cyclosporin family; immunosuppressive and antifungal activities | 88 |
| 163192124 | Cyclosporin C-like | *Purpureocillium* / EtOAc | Insecticidal activity reported for several members of the cyclosporin family; immunosuppressive and antifungal activities | 88 |
| 397 | Indole-3-acetamide | *Purpureocillium* / *n*-BuOH | Plant growth regulator and IAA precursor through the auxin biosynthetic pathway | 89, 90 |

**Supplementary Material 4.** Selected putatively annotated strain-associated compounds and their reported biological activities.
